# Stoichiometry and compositional plasticity of the yeast nuclear pore complex revealed by quantitative fluorescence microscopy

**DOI:** 10.1101/215376

**Authors:** Sasikumar Rajoo, Pascal Vallotton, Evgeny Onischenko, Karsten Weis

**Affiliations:** Institute of Biochemistry, Department of Biology, Swiss Federal Institute of Technology Zurich (ETH Zurich), CH-8093 Zurich, Switzerland; Molecular Life Science PhD Program, Life Science Zurich Graduate School, CH-8057 Zurich, Switzerland

**Author notes:** These authors contributed equally to this work (Sasikumar Rajoo, Pascal Vallotton).

**Keywords:** nuclear pore complex, nucleoporins, stoichiometry, quantitative fluorescence microscopy, NPC composition

## Abstract

The nuclear pore complex (NPC) is an 8-fold symmetrical channel providing selective transport of biomolecules across the nuclear envelope. Each NPC consists of ~30 different nuclear pore proteins (Nups) all present in multiple copies per NPC. Significant progress has recently been made in the characterization of the vertebrate NPC structure, however, because of the estimated size differences between the vertebrate and yeast NPC, it has been unclear whether the NPC architecture is conserved between species. Here, we have developed a quantitative image analysis pipeline, termed Nuclear Rim Intensity Measurement or NuRIM, to precisely determine copy numbers for almost all Nups within native NPCs of budding yeast cells. Our analysis demonstrates that the majority of yeast Nups are present at most in 16 copies per NPC. This reveals a dramatic difference to the stoichiometry determined for the human NPC suggesting that despite a high degree of individual Nup conservation, the yeast and human NPC architecture is significantly different. Furthermore, using NuRIM we examined the effects of mutations on NPC stoichiometry. We demonstrate for two paralog pairs of key scaffold Nups, Nup170/Nup157 and Nup192/Nup188 that their altered expression leads to significant changes in Nup stoichiometry inducing either voids in the NPC structure or substitution of one paralog by the other. Thus, our results not only provide accurate stoichiometry information for the intact yeast NPC but also reveal an intriguing compositional plasticity of the NPC architecture, which may explain how differences in NPC composition could arise in the course of evolution.

**Significance:** The nuclear pore complex (NPC) is one of the largest protein complexes in eukaryotes comprising over 500 nucleoporin subunits. The NPC is essential for transport of biomolecules across the nuclear envelope, however, due to its enormous size, it has been a challenge to characterize its molecular architecture. Herein, we have developed a novel, widely applicable imaging pipeline to determine the absolute nucleoporin abundances in native yeast NPCs. This reveals that the NPC composition dramatically differs between yeast and human despite an overall conservation of individual subunits. We also applied our imaging pipeline to examine yeast mutants revealing a remarkable compositional plasticity of NPCs. Our stoichiometry analyses provide an important resource for the generation of high-resolution structure models of the NPC.

## Introduction

In eukaryotes, chromosomes reside in the nucleus, a dedicated cellular compartment that is delimited by the inner and outer membrane of the nuclear envelope (NE). The transport across this membrane barrier is made possible by nuclear pore complexes (NPCs) - very large channels that span and merge the inner and outer nuclear membranes (1). Despite an estimated mass of 60-120 MDa, NPCs are built from only ~30 different building blocks called nucleoporins, or Nups (2–5). NPCs display an axial eight-fold symmetry (6) such that individual Nups are expected to be present in multiples of 8 copies (i.e. 8, 16, 24, etc.) and each NPC is therefore composed of 500-1000 proteins ultimately accounting for its large mass (7–10).

Significant progress has been made in the last few years in the structural characterization of components of the NPC. X-ray crystallography has provided an inventory of atomic structures for a large number of individual Nups, and also for many Nup complexes that form stable subassemblies (11, 12). For example, the Y complex (or Nup84-complex) comprising a single copy of each Nup84, Seh1, Nup85, Nup120, Sec13, Nup145C, and Nup133, has been extensively characterized both biochemically and structurally (13–15). It is expected that subcomplexes such as the Y complex have the same stoichiometry within the context of intact NPCs as they have in vitro, and thus for the Y complex an abundance ratio of 1:1:1:1:1:1:1 is predicted. Furthermore, several groups have analyzed the overall stoichiometry of Nups within the NPC (4, 7, 8, 16–19). Notably, a combination of mass spectrometry and fluorescent quantification was used to conclude that the human NPC consists of 32 copies of most scaffold Nups, including all members of the Y complex (18).

Additionally, remarkable advances in cryo-electron tomography have allowed for detailed structural characterization of intact NPCs exemplified by current structures for Xenopus and human NPCs at resolutions of ~20 and ~23 Å, respectively (20, 21). The EM maps together with stoichiometry data, structural information, genetic data from yeast and a large body of biochemical interaction studies were recently integrated to produce the first molecular models for the whole human NPC scaffold architecture (22, 23).

Whereas individual Nups and Nup subcomplexes are generally well conserved throughout evolution, it is unclear whether the structural model of the human NPC can be applied to all eukaryotes. For example, the molecular mass of the budding yeast NPC was measured in the 60 MDa range compared to 100-130 MDa for the vertebrate NPC (24–27). Additionally, cryo-electron reconstructions show that the yeast NPC is smaller and flatter in comparison to its vertebrate counterpart (25). Careful examination of Nup stoichiometries will be critical to understand the basis of such evolutionary differences. Moreover, reliable copy number information for every Nup is essential to produce faithful 3D models of the NPC structure as stoichiometry together with symmetry information underlies the entire process of fitting the electron density of individual subunits into experimental electron microscopy maps (28).

In this study, we have used a library of yeast strains expressing endogenously GFP-tagged Nups to accurately determine nucleoporin copy numbers within the intact NPC. To this end, we have developed a high-throughput automated fluorescence imaging technique, that we termed Nuclear Rim Intensity Measurement or NuRIM. This allowed us to obtain relative Nup stoichiometries based on the precise quantification of fluorescence intensities at the NE. Comparison of the intensity of selected Nups at the single NPC level with well-characterized fluorescence intensity standards allowed us to obtain the absolute copy numbers for almost all Nups in yeast. Remarkably, our results show that most of the nucleoporins in yeast are present in only half the copy number as compared to their mammalian counterparts (22, 23), thus identifying a striking difference in the composition of yeast and vertebrate NPCs. Furthermore, by exploiting our workflow, we found that manipulation of Nup expression levels led to the formation of functional NPCs with significantly altered stoichiometries revealing an astonishing plasticity of NPC architecture.

## Results

### NuRIM: a quantitative live cell imaging assay to determine the relative stoichiometry of GFP-tagged nucleoporins within intact NPCs

Relative abundances (i.e. stoichiometries) of Nups were comprehensively evaluated both in vertebrate cells and in budding yeast using bulk biochemical methods such as mass spectrometry (18) or quantitative SDS PAGE (4). These methods rely on biochemical purifications where it is difficult to ensure and assess the integrity of the NPC preparations. By contrast, fluorescence imaging approaches have the potential to determine Nup abundances within intact NPCs in living cells. If the tags do not interfere with NPC incorporation, comparisons of average intensities over many NPCs within many NEs expressing different GFP-tagged Nups should provide very accurate relative abundance information. For example, a Nup present at twice the copy number of another should yield an average intensity value at the NE differing by a factor of two (see Appendix 1).

We therefore set out to develop a live cell fluorescence imaging workflow to measure fluorescence signal intensities of endogenously labeled Nups specifically at the NE. To implement it, we created a library of strains endogenously expressing C-terminally tagged Nups with yeast-optimized enhanced green fluorescent protein (yEGFP) preceded by a short flexible linker. To ensure that labeling does not interfere with Nup function and NPC incorporation, the growth rates of all labeled stains were compared to wild-type cells (Fig. S1A) and the Nup-yEGFP localization together with the cellular morphology of the tagged strains were carefully examined. In total we were able to successfully tag 26 different Nups that passed our stringent selection criteria.

To accurately quantify Nup-yEGFP intensities at the NE in an unbiased manner, we developed an automated image processing pipeline (Nuclear Rim Intensity Measurement, or NuRIM). The NE was marked using the ER marker dsRed-HDEL. Yeast cells co-expressing various yEGFP-tagged Nups together with dsRed-HDEL were grown in parallel under the same conditions, and imaged side-by-side in an automated manner using identical microscope settings, both in bright field and for the dsRed and GFP channels (Fig. 1A). Bright field images were used to avoid highly crowded areas (Fig. 1A i and ii), while the dsRed channel was used to produce binary masks outlining the NE contours by an algorithm based on Laplacian edge detection (29) (Fig. 1A iii and iv, Appendix 2). These masks closely overlapped spatially with the Nup-yEGFP distribution, and their medial line (morphological skeleton) was used to compute corresponding background-corrected pixel intensities from the yEGFP channel (30) (Fig. 1A v and vi). These values were averaged for each cell, image and yeast strain over thousands of nuclei to produce a final population average intensity value. Appendices 1 and 2 provide a more systematic description of the rationale and the assumptions that underlie NuRIM.

**Fig. 1.**
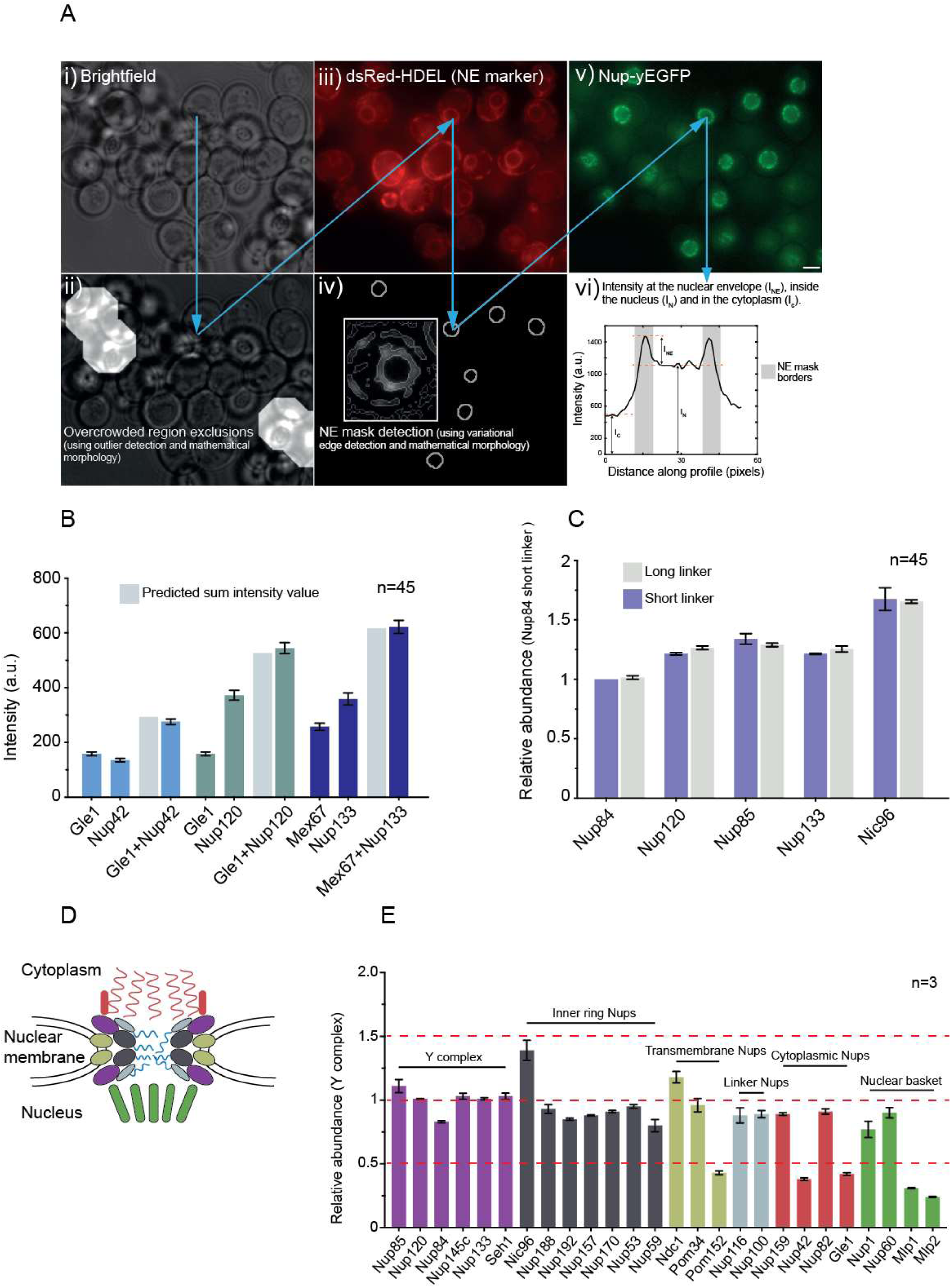
Analysis of the relative nuclear pore complex stoichiometry. (A) Outline of the NuRIM (Nuclear Rim Intensity Measurement) technique. (i-iv) Cells co-expressing dsRed-HDEL and various GFP-tagged Nups were imaged sequentially in bright field, dsRed and GFP channels (i) Bright field channel shows localization of immobilized yeast cells. (ii) Exclusion of the overcrowded regions (light areas) based on the bright field images. (iii) dsRed-HDEL channel shows localization of dsRed-HDEL marking the NEs. (iv) Production of binary NE contour masks based on the dsRed images. (v) GFP channel showing localization of endogenously GFP-tagged Nups. (vi) Background-subtracted Nup-GFP intensities are measured and averaged across thousands of binary NE contours. Scale bars: 2 μm. (B) Linearity of GFP-based intensity measurements. Cells expressing indicated yEGFP-tagged Nups alone or in pairwise combinations were imaged in parallel. Corresponding NE signal intensities were quantified using NuRIM. Grey bars represent corresponding arithmetic sums of the NE signal intensities of singly labeled strains. Mean ± SD; n - number of image frames analyzed. (C) Robustness of GFP-based intensity measurements against micro-environmental variability. Intensity values obtained using NuRIM for the indicated Nups with C-terminal yEGFP tag separated by 6 or 102 amino acid linker sequence. Intensity values were normalized to Nup84-yEGFP containing a short linker. Mean ± SD; n - number of image frames analyzed. (D) Schematic representation of the NPC. Nucleoporin subgroups are depicted as colored cartoons. (E) Relative abundance of yeast Nups. NE intensities for the indicated yEGFP-tagged Nups were quantified using NuRIM and normalized for the average value obtained for the six components of the Y complex. Mean ± SD; n - number of replicates analyzed and are representative of three independent experiments performed on different days.

An important assumption of our approach is that each individual Nup-yEGFP contributes linearly to the total intensity emitted from the NPC. To test this, we quantified the NPC intensities in a set of strains expressing individually labeled Nups, including Gle1, Nup42, Nup120 and Nup133, as well as the NPC-associated nuclear transport factor Mex67, and compared their intensities to the values obtained in double-labeled stains that co-expressed various pairs of these proteins. Within less than 5%, the fluorescence intensities of all tested double-labeled strains matched the values calculated by summing up the intensities of the two corresponding strains expressing a single Nup-yEGFP. This shows that our intensity measurements increase linearly in relation to copy number and can be used to measure relative abundance (Fig. 1B).

Another assumption is that neither crowding nor a different ‘microenvironment’ in the NPC significantly influence fluorescence intensities and affect the accuracy of our measurements. Extreme proximity of EGFPs in oligomers can lead to quenching of fluorescence emission by homo-FRET (31, 32), and tight binding of anti-GFP antibodies to GFP was shown to modify fluorescence intensities (33). While the structural symmetries present in the NPC are expected to prevent such immediate contacts of individual GFP probes, we nevertheless set out to obtain evidence that the microenvironment does not significantly affect our intensity measurements. To this end we introduced a 102-amino-acid-long, flexible linker upstream of yEGFP in some of the strains. Such a long linker should release the yEGFP probes from their microenvironment – thus leading to a potential alteration of the fluorescence intensities. However, when we analyzed the corresponding signal intensities, no significant changes were observed compared to the results obtained with regular linkers (Fig. 1C).

Furthermore, for the correct interpretation of the NuRIM results it is important that the spatial probability distribution (denoted p in Appendix 1) of the various analyzed Nups is identical. However, some Nups might be present in only a subset of NPCs or in additional, non-NPC pools at the NE. To recognize such unusual behavior, we compared the distribution patterns of all Nups in our library using normalized cumulative intensity distributions and the autocorrelation of intensities along the NE. For the majority of Nups, we observed highly similar signatures (Fig. S1B and C). The three outliers, Ndc1, Mlp1 and Mlp2, are consistent with the additional presence of Ndc1 at the spindle pole body (34), and with the previously described absence of both Mlp1 and Mlp2 from NPCs in the nucleolar region (35) (Fig. S1F).

Although we were careful to perform all imaging experiments under very similar conditions, we also tested the robustness of NuRIM against inherent variability in culture density and the density of cells plated on the microscope slide (Fig. S1 D and E). To this end, we varied either of these parameters for one of the strains (Nup84-yEGFP) in the range ~0.5 x to 2 x around the standard values used in the experiments and analyzed the impact on the NuRIM output. Within this range, we did not observe any significant influence on the readout (Fig. S1 D and E). Therefore, our imaging and quantification strategy is resilient against small unavoidable variations in cell culture and imaging conditions.

In summary, we conclude that NuRIM allows for an accurate and unbiased determination of the relative stoichiometry of endogenously labeled Nups within intact NPCs of living cells.

### Determination of the relative nucleoporin abundance

Next, we applied NuRIM to systematically quantify abundances for all Nups in our library (Fig. 1D and E). In order to robustly compare sets of measurements performed on separate days, the average yEGFP intensity of six Y complex components was used for normalization. The intensity readouts were highly reproducible and revealed a striking pattern of abundances (Fig. 1E and Table 1). The vast majority of Nups were present in very similar amounts, displaying values within 15 % of each other (Fig. 1E, horizontal line ‘1’ and Table 1). This includes all members of the Y complex and all major components of the NPC scaffold. One exception was Nic96, which was present in significantly higher amounts, displaying an abundance close to 1.5. In addition, Pom152-GFP, Nup42-GFP, and Gle1-GFP displayed distinctly lower abundance values clustering around 0.5. The three outliers that are not evenly distributed along the NE, Ndc1, Mpl1 and Mlp2 are only included for completeness (Table 1) and need to be viewed with prudence. Accordingly, the majority of Nups are present in the same copy number and in general, Nups can be found in well-defined quanta of ~0.5, 1 or 1.5. Such distinct abundance classes are consistent with the 8-fold symmetrical architecture of the NPC in which Nups are expected to be present in multiple copies of 8 per structure.

**Table 1.** Stoichiometry of nucleoporins within the yeast nuclear pore complex.

| NUPs<sup>*</sup> | Fold ratio<sup>a</sup> | Absolute<sup>b</sup> | 8-Fold<sup>d</sup> |
| --- | --- | --- | --- |
| Nup85 | 1.11 ±0.051 | 18.6 | 16 |
| Nup120 | 1.01 ±0.003 | 16.8 | 16 |
| Nup84 | 0.83 ±0.008 | 13.9 ±2.8<sup>c</sup> | 16 |
| Nup145C | 1.03 ±0.024 | 17.2 | 16 |
| Nup133 | 1.01 ±0.009 | 16.8 | 16 |
| Seh1 | 1.03 ±0.025 | 17.2 | 16 |
| Nic96 | 1.39 ±0.079 | 23.3 | 24 |
| Nup188 | 0.93 ±0.034 | 15.6 | 16 |
| Nup192 | 0.86 ±0.008 | 14.3 | 16 |
| Nup157 | 0.88 ±0.005 | 14.7 | 16 |
| Nup170 | 0.91 ±0.009 | 15.2 | 16 |
| Nup53 | 0.95 ±0.014 | 15.9 | 16 |
| Nup59 | 0.80 ±0.048 | 13.3 | 16 |
| Ndc1* | 1.18 ±0.045 | 19.7 | 16 |
| Pom34 | 0.97 ±0.053 | 16.2 | 16 |
| Pom152 | 0.43 ±0.015 | 7.2 | 8 |
| Nup1 | 0.77 ±0.064 | 12.8 | 16 |
| Nup60 | 0.90 ±0.040 | 15.0 | 16 |
| Nup116 | 0.88 ±0.059 | 14.7 | 16 |
| Nup100 | 0.89 ±0.028 | 14.8 | 16 |
| Nup159 | 0.89 ±0.010 | 14.9 | 16 |
| Nup42 | 0.38 ±0.011 | 6.4 | 8 |
| Nup82 | 0.91 ±0.020 | 15.2 | 16 |
| Gle1 | 0.42 ±0.010 | 7.0 | 8 |
| Mlp1* | 0.31 ±0.004 | 5.2 |  |
| Mlp2* | 0.24 ±0.005 | 3.9 |  |
| Kap95 | 1.34 ±0.013 | 22.4 |  |
| Mex67 | 0.82 ±0.012 | 13.7 |  |

\* Nups that failed one of the controls (see Fig. S1).
<sup>a</sup> obtained by normalizing to the Y complex. Mean ± SD; values are obtained from three independent experiments.
<sup>b</sup> absolute Nup copy numbers based on Nup84.
<sup>c</sup> Mean ± SEM; values are obtained from three independent experiments.
<sup>d</sup> 8-fold rendered values were designated based on absolute Nup copy numbers.

### Determination of absolute copy numbers reveals that most yeast nucleoporins are present in 16 copies per NPC

If the absolute NPC copy number was available for even a single Nup, our systematic ratiometric analysis would allow the determination of absolute copy numbers for all other Nups as well. We thus set out to quantify the total fluorescence intensity of a single NPC in cells expressing one of the fluorescently tagged Nups. Of note, endogenous tagging of Nups principally results in 100% labeling efficiency since all copies of the protein carry the fusion tag (but see also discussion). We took advantage of the observation that single NPCs can occasionally be detected in dividing yeast cells where the part of the NE bridging the mother and daughter nuclei elongates and narrows (Movie S1) (36). These NPCs are more separated from their neighbors allowing us to image and quantify the intensity of individual NPCs in several hundreds of cells using the Diatrack particle tracking software (37). As an absolute intensity reference, we also imaged in parallel, yeast-expressed rotavirus-like VP2 particles that contain exactly 120 copies of yEGFP (38). Furthermore, we recorded the intensity for single molecules of yEGFP adsorbed on glass coverslips (Movie S2). The intensity distributions of single yEGFP, Nup84-yEGFP NPCs and VP2-yEGFP particles were fitted with a normal distribution. Comparison of their means showed that VP2 particles were 116±7 times brighter than single yEGFP molecules yielding an estimated copy number for Nup84-yEGFP of 13.9+-2.8 per NPC (Fig. 2A).

**Fig. 2.**
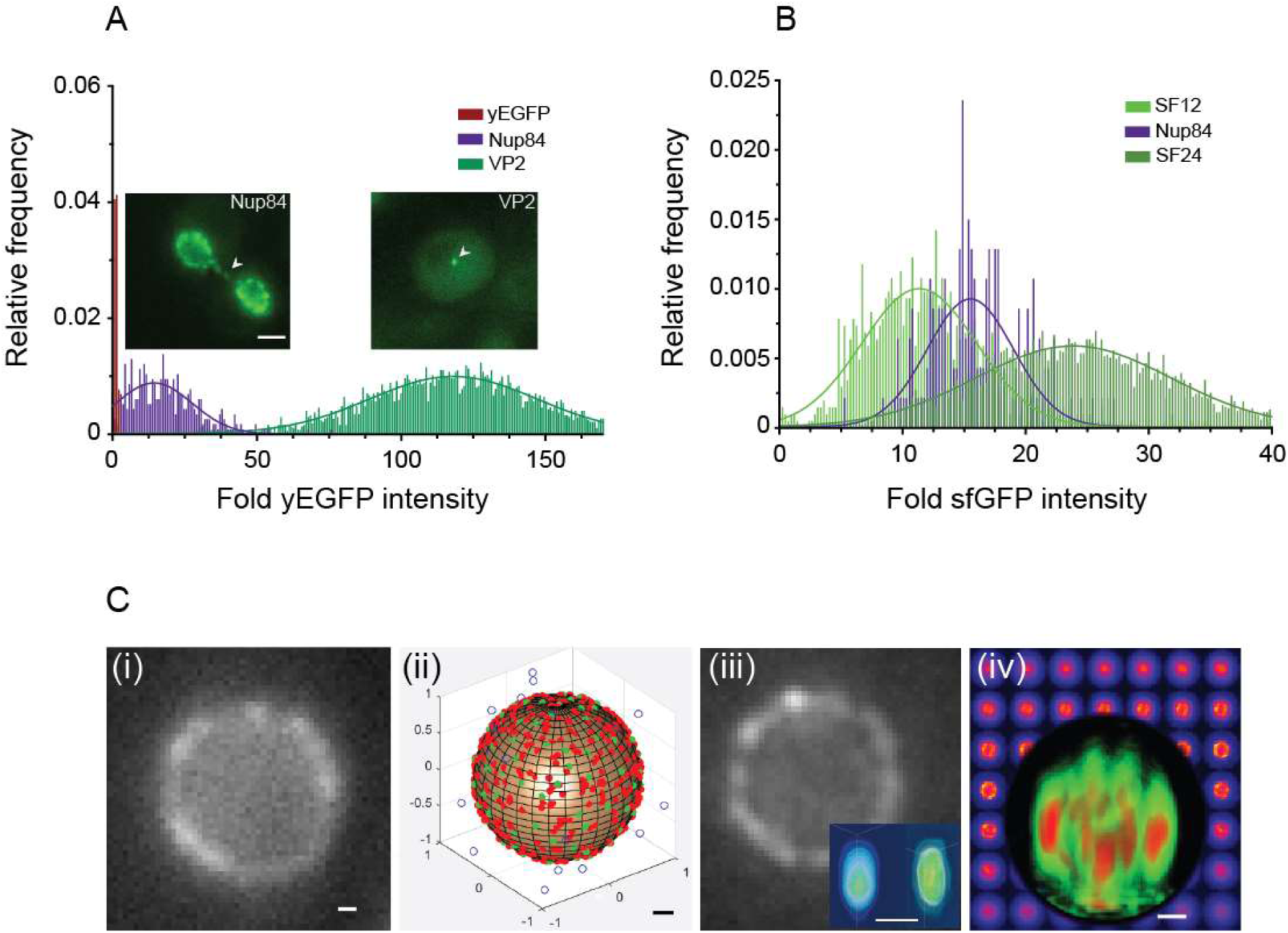
Absolute stoichiometry of yeast nucleoporins. (A) Representative intensity histograms and Gaussian fits (smooth lines) obtained for single yEGFP molecules, single NPCs in regions between separating nuclei in Nup84-yEGFP expressing yeast cells (arrowhead) and for the 120-mer VP2-yEGFP particles produced in yeast (arrowhead). Scale bars: 2 μm. (B) Intensity histograms and fits are displayed similar to (A) for single NPCs in Nup84-sfGFP expressing yeast cells, and purified SF12 and SF24 particles containing 12 and 24 sfGFP molecules, respectively. (C) i) Representative fluorescence microscopy image showing localization of Nup84-yEGFP labeled NPCs at the NE. ii) Computational model of the NE. NPCs are represented as green dots, dsRed-HDEL as red dots, and dim sources simulating background fluorescence as blue dots. iii) Simulated image of the NE produced computationally from the model (ii). Note good resemblance to genuine NEs (compare with i)). Inset compares computed PSF with experimentally acquired PSF. (iv) Representative 3D Rendering of image stack shows heat map of intensities in Nup84-yEGFP cells illustrating high intensity spots or ‘speckles’ (red) produced due to stochastic overlap of the ~120 NPCs. Scale bar represents 200 nm.

To validate this result, we performed an independent analysis using a different set of reference intensity markers: tetrahedral particles that contain precisely 12 or 24 GFP moieties (39). These particles contain ‘superfolder’ GFP (sfGFP), a GFP variant with fast folding kinetics (40), and we therefore endogenously tagged several Nups including Gle1, Nup84, Nup120 and Nic96 also with sfGFP. As expected, tagging with sfGFP did not lead to any significant changes in the relative abundance readouts as compared to yEGFP (Fig. S2A). Using the same methodology as before, we then recorded in parallel the absolute intensity distributions for 12x sfGFP and 24x sfGFP tetrahedral particles purified from E. coli, and for the Nup84-sfGFP and Nup120-sfGFP NPCs in yeast. Based on the comparison of the means of the distributions we determined absolute NPC copy numbers of 14.7+-2.5 for Nup84 and 15.8+-2.3 for Nup120 (Fig. 2B and S2B).

In accordance with these results, we assigned absolute stoichiometries for all Nups measured in Table 1. We conclude that the yeast NPC contains very close to 16 copies for most Nups (Table 1). Most notably, these include the majority of scaffold Nups constituting the framework of the NPC: the Y complex members (Nup133, Nup120, Nup85, Nup84, Nup145C, Seh1) and the inner ring Nups (Nup188, Nup192, Nup157, Nup170). These numbers are in stark contrast to the human NPC where corresponding proteins were reported to be generally twice as abundant, including 32 copies of the Y complex (16, 21–23). The astonishing differences in NPC composition between yeast and human prompted us to develop another independent method to examine NPC stoichiometry. The characteristic intensity spots (i.e. speckles) that are observed by fluorescence microscopy in the yeast interphase NE are produced by randomly overlapping, diffraction-limited images of several closely positioned NPCs (Fig. 2C iv). Precise determination of a characteristic ‘NPC clustering’ factor within such spots would allow us to estimate an absolute stoichiometry by comparing their intensities with established external fluorescence standards. We thus developed a 3D mathematical model of the NE based on NPC density data derived from 3D electron microscopy reconstructions (36) and on classical image formation theory (Appendix 2). This model allowed us to estimate the characteristic ‘NPC clustering factor’ predicting that on average 3.7 NPCs are present in such NPC intensity spots (Appendix 2). 3D stacks acquired for Nup84-yEGFP and VP2-yEGFP expressing cells imaged under identical conditions were then used as inputs to quantify absolute signal intensities. By comparing the intensity distributions of the NPCs spots and the VP2 particles, we computed a copy number of 13.8+-4 for Nup84 per NPC. This result is fully consistent with - and confirms our measurements based on single NPC intensities (Fig. 2C).

Altogether, our results demonstrate that on average no more than 16 copies of Nup84 are present in the yeast NPC. Based on our relative abundance measurements, most Nups within the NPC of budding yeast therefore exist in 16 copies. This finding is consistent with molecular mass estimates for the yeast NPC, which is significantly smaller compared to its vertebrate counterpart (24, 25). Furthermore, this suggests that there are considerable differences in the architectural organization of the yeast and vertebrate NPCs. From an evolutionary perspective, this implies that a macromolecular machine with conserved function and constructed from conserved subunits can adopt fundamentally distinct structural layouts in different species.

### Altering nucleoporin expression leads to NPCs with different stoichiometries

Our automated imaging pipeline can also be applied to mutant yeast strains in which the composition of the NPC is altered. This paves the way for investigating a wide range of questions regarding the NPC architecture that would have otherwise been difficult to address. For example, in recently proposed models of the human NPC (22, 23), Nup155 is an abundant Nup present in 48 copies playing a critical role in linking the Y complex to the NPC core inner ring and nuclear membrane. Yet, despite this central structural role, in yeast its function is split between two non-essential and functionally redundant paralogous genes NUP170 and NUP157 that have separated due to whole genome duplication ~100 million years ago (41–43). Of note, each of these Nups is present in only ~16 copies in the yeast NPC (Table 1). To study this further, we utilized NuRIM to investigate whether Nup157 can replace Nup170 when the NUP170 gene is deleted. We hypothesized that if substitution was to occur, we should see an increase in the intensity due to additional copies of Nup157 now at the pore. However, in NUP157-yEGFP cells the deletion of NUP170 did not cause any increase in Nup157-yEGFP intensity (Fig. 3A and 3B), indicating that sites that are normally occupied by Nup170 remain empty in its absence. To rule out the possibility that protein expression is a limiting factor for substitution, we constructed a NUP157-yEGFP nup170Δ strain that contained an extra copy of NUP157-yEGFP inserted into the URA3 locus thus doubling the production of this protein. This however, also did not cause any significant increase in its abundance at the NPC. In contrast and as expected, the re-expression of NUP170 by insertion of a NUP170-yEGFP gene into the URA3 locus produced twice as bright NPCs (Fig. 3A and 3B). We conclude that Nup157-yEGFP is not able to substitute for the loss of Nup170 indicating that yeast NPCs with only ~ 16 copies of Nup157 are fully functional.

**Fig. 3.**
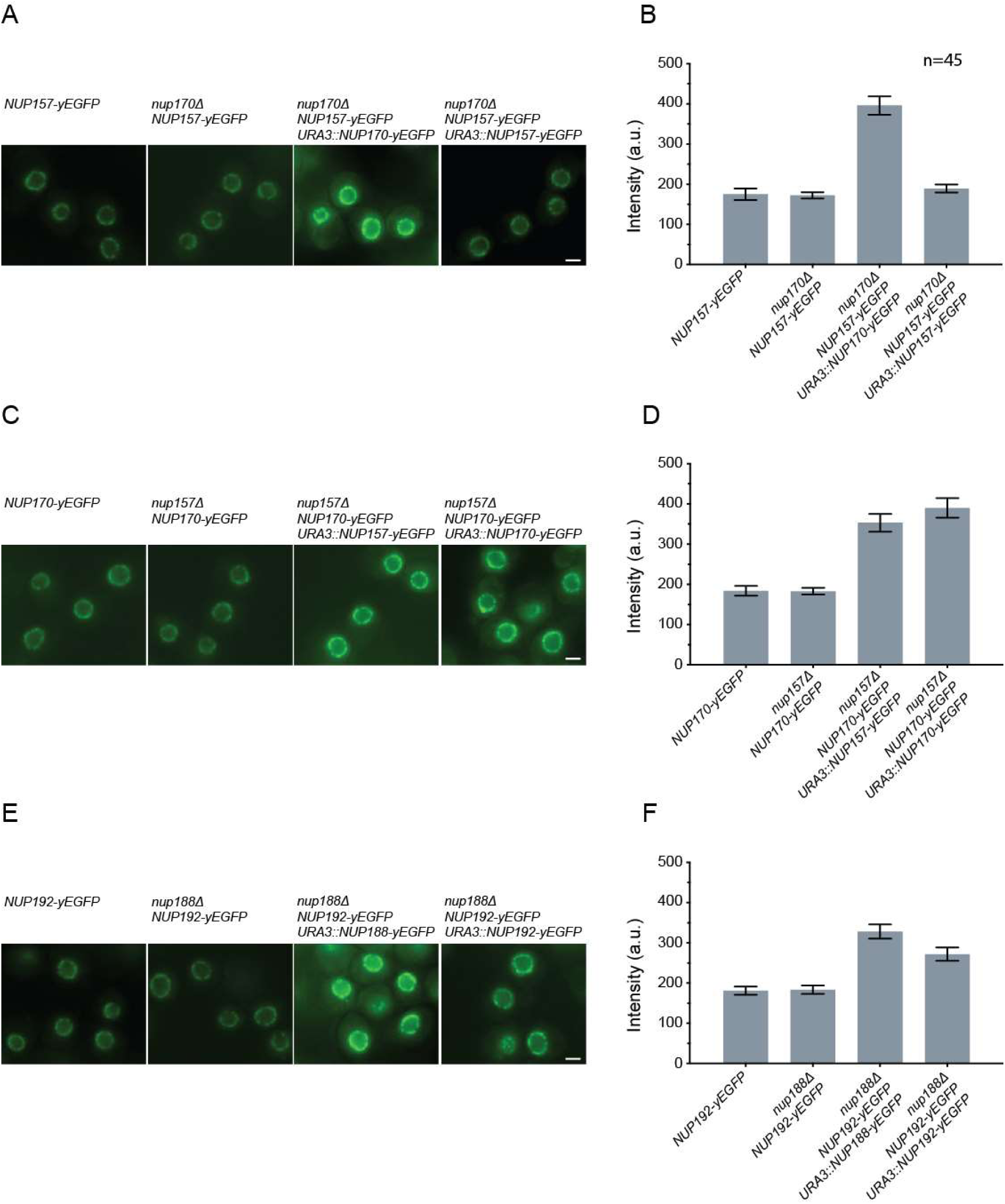
Analysis of nucleoporin stoichiometry in yeast mutants. (A, C and E) Representative images illustrating Nup-yEGFP localization in actively growing yeast cells with specified genotypes. The corresponding NE intensities quantified using NuRIM are shown in B, D and F respectively. Mean ± SD; n - number of image frames analyzed for B, D and F. Scale bars: 2 μm.

Gene duplications supply the raw material for the emergence of new functions through the forces of mutation and natural selection (44). We therefore also wanted to test the inverse scenario i.e. whether Nup170 could substitute for the loss of Nup157 at the NPC. As in the mirror strain, we did not observe any increase in abundance of Nup170-yEGFP upon deletion of NUP157 (Fig. 3C and 3D). However in this case, the introduction of an additional copy of NUP170-yEGFP led to a 2-fold increase in Nup170 abundance at the NPC (Fig. 3C and 3D). This reveals an interesting functional asymmetry between these two paralogs and shows that manipulating gene expression levels can change the NPC stoichiometry producing an NPC with ~32 copies of Nup170.

Motivated by these findings, we also investigated another paralog pair, comprised of Nup188 and Nup192. These two paralogs presumably originated from an ancient gene duplication event - about 800 million years ago (45). Since the gene encoding for Nup192 is essential we restricted our analysis to the deletion of NUP188. Similar to the results with Nup170 and Nup157, no intensity increase was observed in nup188Δ cells in the presence of endogeneous levels of Nup192-yEGFP (Fig. 3E and 3F). However, once an extra copy of NUP192-yEGFP was introduced, an increase in NPC abundance of more than 50% was detected (Fig. 3E and 3F). This suggests that Nup192 is capable of occupying the empty Nup188 sites but it is unable to do so entirely. Taken together, these results identify intriguing redundancies between Nup paralogs and reveal a remarkable compositional plasticity of the NPC.

### Discussion and conclusion

The inventory of Nups that make up the NPC has been biochemically characterized in several organisms from diverse branches of the eukaryotic evolutionary tree (2–5). Whereas individual components of this inventory are largely well conserved, it has been unclear to which extent this is also true for the architecture of the entire NPC. To understand how the vast number of NPC building blocks come together to produce a functional structure requires various sources of high quality data including, precise information about the copy number for each Nup subunit (28). In this study, we have developed a strategy based on live cell imaging to accurately determine the absolute stoichiometry for the majority of Nups within the intact budding yeast NPC, and applied this approach to begin to probe the principles underlying the structural organization of the NPC using nucleoporin mutants. Our analyses revealed striking differences between yeast and human NPCs, and exposed an inherent compositional plasticity of the NPC architecture.

In budding yeast, NPCs are densely packed within the NE (36) preventing resolution of individual structures by light microscopy. However, NPCs containing twice as many copies of GFP-labeled Nup when imaged identically will nevertheless give rise to NEs that are twice as bright. Our approach to determine the stoichiometry of the NPC was based on this principle and relied on precise measurement of fluorescence intensities of endogenously GFP-tagged Nups at the NE to evaluate their relative abundance (46) (Fig. 1E). These measurements were automated and subjected to multiple layers of quality control to ensure that the results (a) are linearly dependent on the Nup copy number, (b) are highly reproducible between multiple measurements of thousands of nuclei, (c) are robust to small variations in experimental conditions, and (d) are not influenced by variations in the GFP micro-environment for the various tagged Nups (Fig. 1 and S1). The relative stoichiometries obtained with this approach are consistent with an 8-fold symmetrical NPC organization, showing that all 23 Nups that could be reliably measured cluster around three distinct quanta of abundance (0.5, 1.0 and 1.5). In contrast, NTRs that are dynamically associated with the NPC do not show such a clear discretization in their abundance (Fig. 1E and Table 1). In our analysis, we found that three Nups, Ndc1, Mlp1 and Mlp2 displayed an atypical distribution pattern in the NE, and also these three outliers deviated form the discrete abundance pattern of other Nups (Fig. 1E and Table 1). The numerical value for the abundance of Ndc1, which is slightly higher compared to the Y complex members is likely explained by the additional presence of Ndc1 at the spindle pole body (34), whereas the exceptionally low values for Mlp1 and Mlp2 is consistent with their absence from a subset of NPCs (Fig. 1E, S3F and Table 1).

Overall, when compared to a previous comprehensive stoichiometry analysis in yeast by Rout and colleagues (4), our results are in good agreement for symmetric core Nups showing approximately similar relative abundances, but differ significantly for several peripheral Nups, which we find in higher amounts (Fig. 1E and Table 1). This discrepancy could be explained by a partial loss of peripheral Nups that might be difficult to avoid during the biochemical purification of the NEs used in this prior work. Interestingly, distinct copy numbers of yeast Nups were previously also reported using step bleaching of single NPCs with an intense laser spot at the NE (17). However, since the punctae seen with fluorescently labeled NPCs in regular regions of yeast interphase NE mostly do not correspond to single structures but to the randomly overlapping images of at least a few of them, it is unclear to us how imaging of single NPCs was guaranteed in this study (Fig. 2C) nor how NPCs were immobilized over the time scale of several seconds needed for recording photo-bleaching curves.

In mammalian cells, the typical distance between neighboring NPCs in the NE lies generally above the Abbe resolution limit (10), making it possible to characterize fluorescence signals from NPCs one at a time with optical microscopy (47). As pointed out, this is not the case in interphase budding yeast cells, but during cell division the NE between the dividing mother and daughter nuclei extends and narrows which distributes individual NPCs in this region sufficiently to resolve them by light microscopy (36). We used this phenomenon to quantify absolute Nup copy numbers within individual NPCs by precise brightness measurements of GFP-tagged yeast strains and comparing those to a set of fluorescence standards with a defined number of identical GFP molecules. The copy number values that we determined for two different Y complex members, Nup84 and Nup120, were close to 16 copies per NPC. Very similar copy numbers were obtained with different fluorescent tags and fluorescent standards (Fig. 2 and S2). Moreover, the conclusion that there are 16 copies of Nup84 per NPC was further confirmed by a 3D modeling approach that was based on the measured NPC density (36) and absolute intensity measurements of NPC clusters in interphase NEs (Fig. 2C). Potentially, our estimation of absolute copy numbers could be influenced by GFP maturation time, resulting in different fractions of fluorescent GFP molecules in NPCs or in our standards. While this could explain why we find copy numbers for Y complex components that are slightly below 16, the contribution of GFP maturation time is not a significant source of error since similar values were obtained using both yEGFP and sfGFP, which have very different maturation kinetics (48).

Based on our relative abundance and absolute copy number measurements, the vast majority of yeast Nups are present in no more than 16 copies per NPC (Fig. 1E and Table 1). This fits well with prior total mass estimates of 45-70 MD for the yeast NPC (24, 25), but reveals a remarkable difference in structural organization compared to the mammalian NPC: many yeast structural core Nups are present in ~50% lower amounts per NPC compared to their mammalian counterparts (18). One particularly prominent difference is the copy number of the Y complex (also Nup107-or Nup84-complex), an evolutionary conserved architectural element of the NPC accounting for ~1/3 of its total mass, for which we find 16 copies in yeast whereas 32 copies were assigned for the mammalian NPC. Since structural analyses of the mammalian NPC show a peculiar arrangement of the Y complex in two concentric rings on each side of the NPC (16, 21-23), it is likely that the yeast NPC contains only one such ring on the nuclear and cytoplasmic side, respectively. Another obvious difference concerns the vertebrate Nup155 (Nup170 and Nup157 in yeast), which forms multiple important connections, including links to the Y complex, to other core Nups and to the nuclear membrane (49). It is present in 48 copies in the mammalian NPC (22, 23), whereas yeast contains only 16 copies for each of its two paralogs, Nup170 and Nup157, amounting to a total maximum of 32 copies (Fig. 1E and Table 1). Similarly, the evolutionary conserved paralog pair Nup188 and Nup205 (Nup188 and Nup192 in yeast) is present in 16 and 32 copies in mammalian NPC (23), while the yeast NPC contains only 16 copies of each (Fig. 1E and Table 1), suggesting that these conserved paralogs also differ in their abundances between the two species.

These intriguing differences raise the question of how the same set of conserved Nups can produce assemblies that are so diverse in their architectural layout and yet remain functionally equivalent. To begin addressing this, we employed our quantitative approach to analyze the stoichiometry of NPCs in cells that had altered expression of several Nups. First, this showcased that the NPC scaffold is remarkably robust and has multiple inbuilt redundancies. For example, the presence of half the copy number of the Nup170/157 paralog pair is sufficient for viability indicating that NPCs that are built in the absence of either member contain voids in their structure, yet remain fully functional (Fig. 4 B and C). Secondly, our analyses revealed an astonishing compositional flexibility of the NPC architecture. Overexpression of one of the paralogs, Nup170, in the absence of Nup157 produced NPCs that contained 32 copies of solely Nup170 (Fig. 4C). Even more surprisingly, similar changes in NPC stoichiometry were observed for the Nup192/Nup188 paralog pair anciently separated in the eukaryotic evolutionary tree (Fig. 4D). Together, this indicates that the composition of the NPC has inherent plasticity and can be altered by varying the expression levels of its subunits. This could explain how variations, initially driven by alterations in expression levels could result in stable changes in the overall layout of the NPC structure over the course of evolution. Other sources of change may include the acquisition of new NPC components. For instance, the outer Y complex ring accounting for ~25% difference in the copy number between the human and yeast NPC, is attached to the human NPC via Nup358, a Nup that is not present at all in yeast (21). This compositional plasticity suggests that the organization of the NPC is modular - following a “Lego-like” principle, where a set of common components can be utilized to form distinct structures with increasing complexity (Fig. 4). In support of this view, several highly structured NPC subunits are chiefly connected by highly flexible, disordered motifs, such as short linear motifs or FG repeats to ensure NPC stability (49, 50). This modularity could also be a reflection of the shared ancestry of core components of the NPC and COP-coats, in which changes in coatomer copy number drive structural diversity (51, 52). Moreover, this raises the intriguing question whether NPCs even within a single cell might display variability and whether all NPCs in a cell have an identical composition or architecture.

**Fig. 4.**
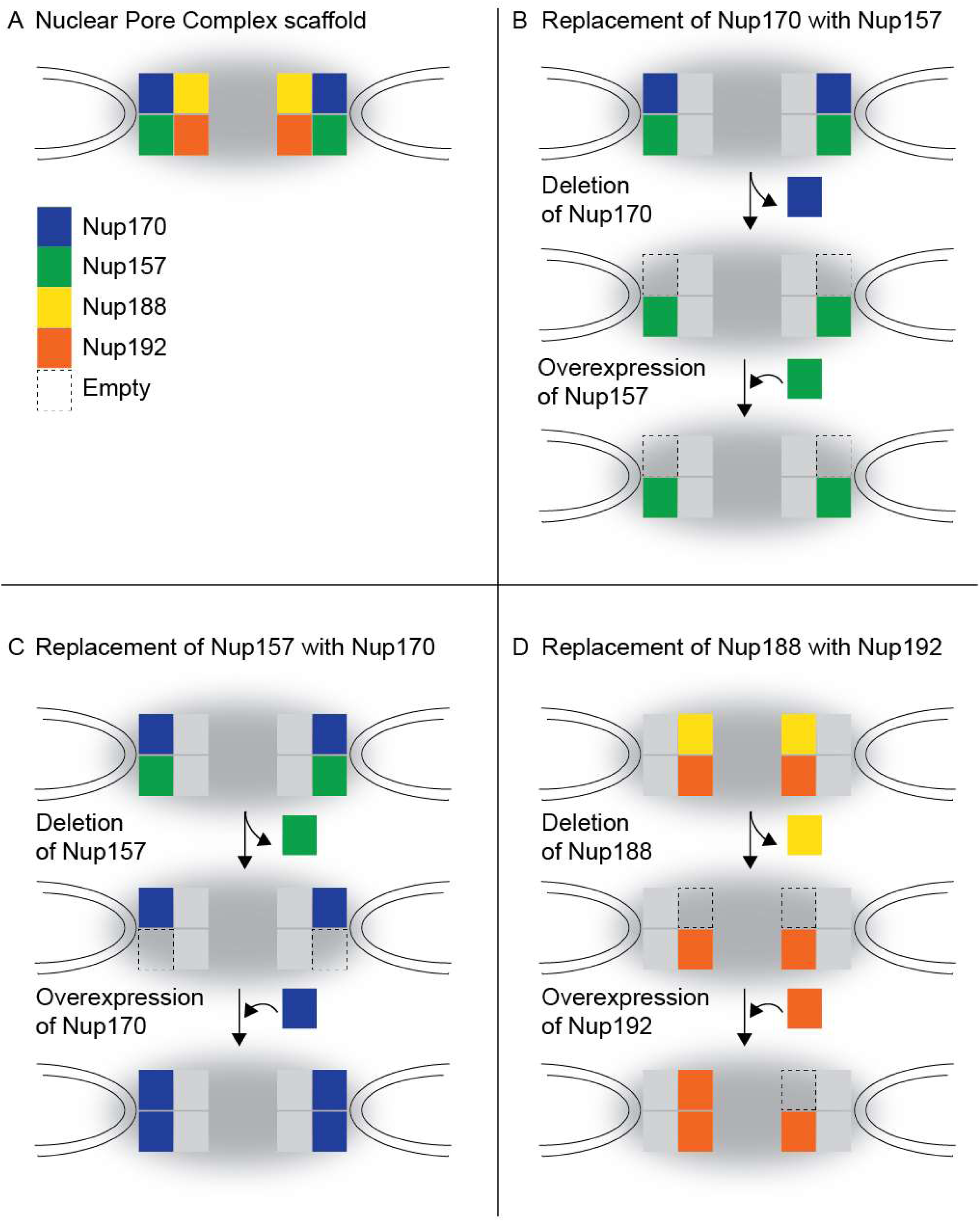
Modular organization of the NPC. (A) Schematic of the NPC scaffold. (B-D) Illustration depicting the experimental outcomes shown in Fig. 3.

In conclusion, we have developed a robust method to accurately quantify both relative and absolute copy numbers for Nups within the yeast NPC. The results of our Nup stoichiometry measurements provide an important resource for the generation of high-resolution models of the NPC. Furthermore, our approach can be easily implemented and applied to other cell biological questions; for example, it may be used to study NPC biogenesis, to analyze NPCs from other species or even to examine other multisubunit complexes in the NE.

## Materials and methods

### Yeast strains and plasmids

The growth media, Saccharomyces cerevisiae strains and plasmids used in this study were produced using standard yeast and molecular cloning protocols, and are listed in Table S1 and S2.

### Growth tests

Yeast strains were grown in synthetic complete medium containing 2% dextrose (SCD). OD600 measurements were acquired over 24 h using CLARIOstar automated plate reader (BMG Labtech) at 30°C in 24-well plates (Nunc) from overnight pre-cultures diluted 1:125 in SCD. Log-phase duplication times (Fig. S1A) were determined from the OD600 reads as average duplication time tdup over the duration of log-phase, a period when tdup reaches minimum and stays approximately constant. tdup was calculated as Δt/Log2(ODt+ Δt/ODt), where ODt+ Δt and ODt are OD600 values at consequent reads and Δt time between the reads.

### Purification of tetrahedral particles

The tetrahedral particles (both SF12 and SF24) were expressed and purified from E. coli (BL21 DE3) as previously described in (53).

### Fluorescence microscopy techniques

Yeast cells endogenously expressing yEGFP tagged Nups were grown to log-phase and imaged in ConA-coated glass bottom plates (MatriPlate) using an inverted epifluoresence Ti microscope (Nikon) equipped with a Spectra X LED light source (Lumencore) and Flash 4.0 scMOS camera (Hamamatsu) with the 100X Plan-Apo VC objective NA 1.4 (Nikon). The NIS Elements software (Nikon) was used for image acquisition. All yeast strains used for the quantitative analysis of Nup-yEGFP intensity also expressed dsRed-HDEL. Image segmentation and GFP signal quantification were performed as outlined in Appendix 1.

Imaging of the tetrahedral particles was done by mounting them on agarose pads for microscopy as previously described in (54).

Image quantification and modelling was conducted as described in Supplementary information (Appendices 1 and 2).

## Acknowledgements

We are grateful to E. Dultz, A. Kralt and other Weis lab members for helpful discussions and suggestions and also to B. Hofland for preparation of various experimental materials. We are indebted to D. Baker (UW) for providing the tetrahedral constructs, and we thank J. Cohent (INRA) for the VP2 construct and C. Weber for preparing a VP2-expressing yeast strain. This work was supported by a grant of the Swiss National Science Foundation to KW (SNF 159731).

## Author contributions

SR performed yeast experiments and PV developed the analysis software. SR, PV, EO and KW participated in experimental design. SR, PV, EO and KW wrote the paper.

## 1) Supplementary Methods

### Appendix 1

#### Rationale for stoichiometric method and underlying assumptions

The principle of the method is best appreciated by considering the probability p(r) of finding a NPC at position r, the light intensity distribution in the image plane I(r), the molecular brightness per yEGFP label: B, the point spread function P(r), and the copy number of a particular yEGFP-tagged Nup: N. The average intensity may then be expressed by:

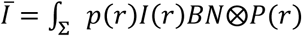

Trivially, N may be taken out of this sum. Therefore, provided that the underlying probabilistic distributions are identical for two different Nup-yEGFP strains, the ratio of their average intensities will estimate precisely the ratio of their copy number.

While conveying the essence of the method, the last expression is not quite satisfactory, as it does not make explicit the averaging process that takes place. For a number K of NPCs at position ri, i=1…K, let again B be the brightness of a single yEGFP tag, N the copy number for the relevant Nup, and E(r) a factor that models inhomogeneous excitation in the field of view. In the absence of diffraction, the emitted intensity distribution is given by:

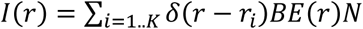

The effect of imaging by diffraction limited optics is captured by a point spread function P(r) via the convolution: 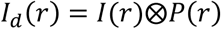.

The effect of sampling by an imaging sensor is modeled by integration over the domain of single pixels:

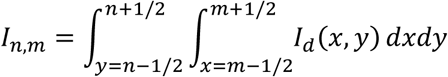

The process of fluorescence intensity averaging over a discrete mask obtained by segmentation of the dsRed-HDEL channel image of size n x m is summarized by a characteristic function χ equal to 1 in the mask, and 0 outside:

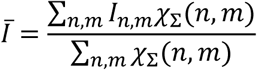

In order to describe the statistical averaging that is taking place (whereby every image is a stochastic realization), one may introduce the probability distribution 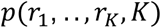) that fully describes the probability of finding K NPC’s at positions 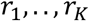.

One may then compute the expected values of any random variable under 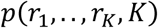. For example, that of 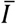

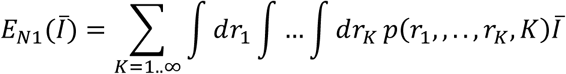

Consider now two different yEGFP tagged Nups with their copy number N1 and N2 and characterized otherwise by identical statistical properties. In particular the labeling by yEGFP is assumed not to deform nuclei in different ways for different mutants. The masks χ_Σ_ may then be assumed to depend on 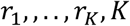 in an identical manner for both mutant (note that the masks do not depend on the copy numbers because dsRed-HDEL is used to compute them instead of the yEGFP tagged Nups themselves). Then for any arbitrary value of 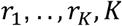, the integrand of *E*_*N1*_ and *E*_*N2*_ will be in a N1:N2 ratio, as can be verified directly by substituting the expression of 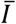 into the last equation. As a result, *E*_*N1*_ and *E*_*N2*_ will also be in a N1:N2 ratio.

It is important to recognize the limitations of our method summarized by these expressions. For example, if the unexpected consequence of introducing yEGFP for a particular Nup were to considerably increase the nuclear size, copy numbers for that particular Nup would probably be underestimated. We therefore introduced several strict controls to flag exceptional Nup-yEGFP phenotypes for which our method may possibly produce biased estimates. First, the visual appearance of various yEGFP tagged Nups were checked for unusual phenotypes. Second, nuclear radii were measured. Third, the cumulative distribution of intensities among all pixels of the nuclear envelope was plotted as a detector for outlier intensity clustering behavior. Fourth, the intensity auto correlation function along the nuclear envelope was computed. Finally, we acquired growth curves and measured doubling times for all GFP tagged Nup constructs (Fig. S1A). Among all the Nups, Mlp1, Mlp2 and Ndc1 failed at least one of our tests while all the other Nups passed them.

#### Image analysis pipeline for yeast NE segmentation

Our methodology for tracing the cell outline was briefly described in the main text. In order to find the boundaries of the NE marked by dsRed-HDEL, we use a Laplacian edge detector.

As a zero level set, this detector ensures that the contours that are detected are both closed and optimal in a variational sense (1). We then analyze the properties of the connected regions enclosed by the closed contours and retain exclusively those regions (i.e. masks) that are compatible with NEs. In particular, we limit the mask selection to those regions only that are thinner than 10 pixels (i.e. 0.6 μm) and that present an average intensity significantly higher than the background as determined using quantile intensity statistics. Further, we avoid quantifying regions of the image where cells exceptionally pile up on each other by analyzing the bright field images and computing exclusion masks (again using quantile statistics). It is important to select a NE marker that is different from the Nups that one aims to compare: by selecting dsRed-HDEL, even potentially low-abundance Nups may still be quantified reliably even if segmenting the NE based on their fluorescence signal might not be possible. Our Matlab code can be found in the supplementary information.

### Appendix 2

#### Generative modeling of NPCs in the nuclear envelope and simulated imaging enabling determination of average number of NPC’s per NE speckle

In order to illustrate and help with the interpretation of the results of our fluorescence intensity measurements, we built a generative 3D model of the nuclear envelope populated by NPCs (Fig. 2C). Nuclear envelopes in yeast are approximately spherical with a mean average radius of 1.0 μm and they were modeled as spheres of that size (2, 3). NPCs were distributed randomly on the surface of the sphere (4). Further, it was observed by Winey et al. that the nearest neighbor distribution between NPCs displays an exclusion zone of approximately 130 nm, presumably due to steric effects between NPC’s (3). To replicate the cumulative nearest distance distributions we used ancestral sampling - rejecting new NPC instances that would lie closer than this cutoff of 130 nm, until a total of 120 NPCs were produced. The value of 120 NPC per cell on average was also obtained from the work of Winey et al.

To produce sample images of NEs, simulated NEs were blurred using a point spread function (PSF) computed using the code from Pankajakshan et al: (5). A computed PSF was used rather than an experimental PSF because it is not trivial to obtain a fully satisfactory PSF experimentally (However, see Fig. 2C). The computed PSF was convoluted with our simulated data using the speed-optimized function “convolution3D_FFTdomain.m” contributed by Christopher Coello (6). This process yielded realistic images of NEs, such that it was difficult to distinguish synthetic images from genuine ones (compare Fig. 2C i and iii)). In order to take into account the fact that nuclei may lie anywhere within the cell, a single slice was selected randomly and uniformly from the 10 slices closest to the brightest one. A simulated intensity standard was introduced in the volume at a random position to simulate the VP2 intensity standard.

Our model was used in particular to determine by which factor on average the intensity of speckles in the NE were brighter than those of isolated NPCs, far from other any NPCs in our experimental conditions. As mentioned in the text, this clustering factor was determined to be 3.7 (Fig. 2C iv). To limit the effect of photo bleaching on our results, experimental data were acquired using a spacing of 0.3 micrometer between Z-slices with total 11 slices spanning the cell volume almost entirely. The parameters used in the simulations exactly reflected those used to acquire experimental Z-stacks of NEs and VP2 particles. This particularly included the minimal distance between local intensity maxima, the requirements on their contrast and the image filtering kernel width - which constitute a set of parameters critical for identifying NE speckles and measuring their average intensity.

The model was also used to assess to what extent violations of our assumptions regarding e.g. the number of the NPCs at the NE, or the nuclear diameter, could influence the clustering value and therefore to estimate the uncertainty obtained for absolute copy numbers based on this methodology. Finally, the model allowed us to follow step by step how the properties of linearity and spatial invariance of classical optical systems justify a ratiometric approach to determine Nup stoichiometry in yeast.

## 2) Supplementary Figures

**Fig. S1.**
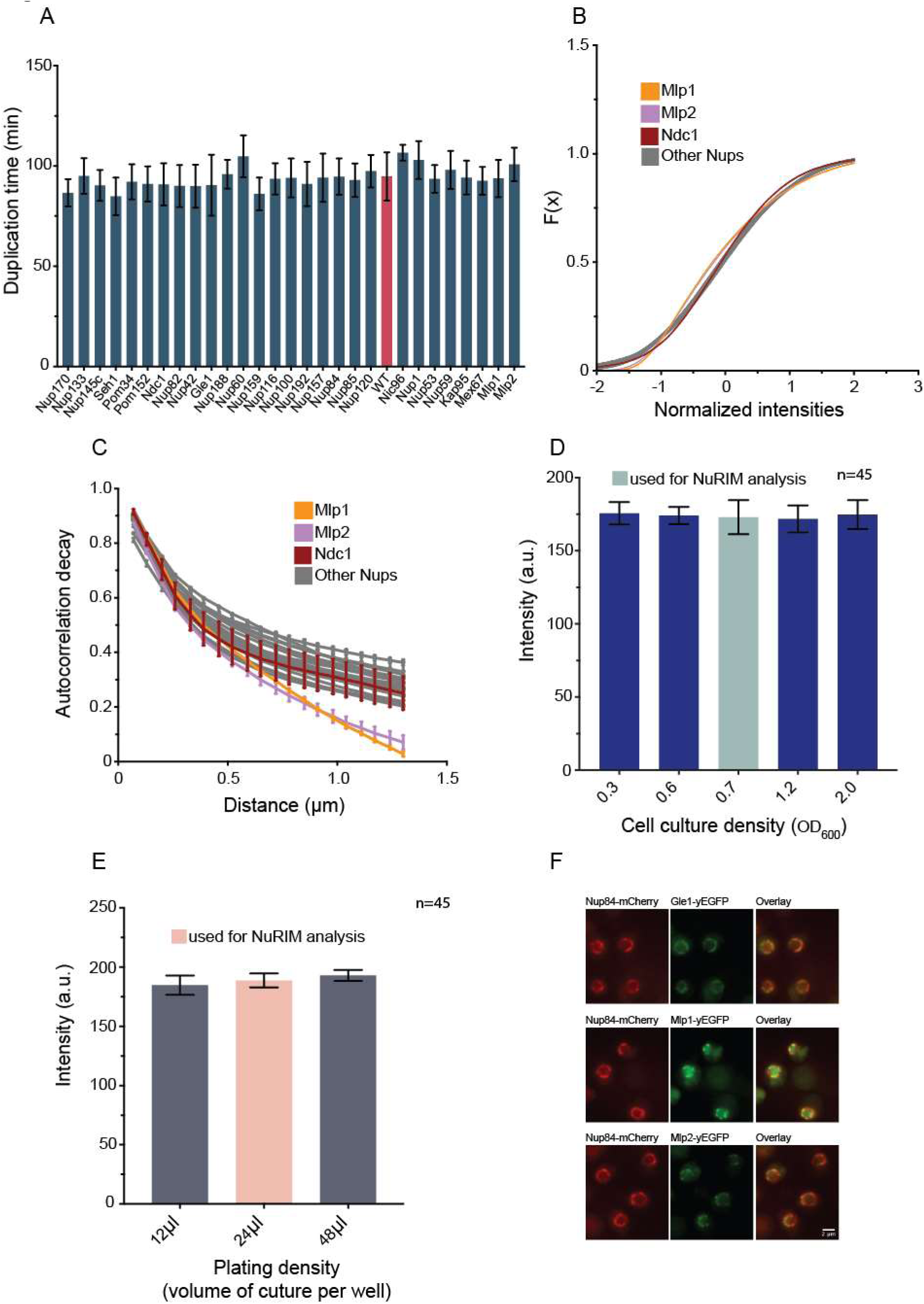
Controls to flag exceptional Nup-yEGFP behavior and analysis of the robustness of NuRIM. (A) Log phase doubling rates determined for the indicated Nup-yEGFP strains grown in SCD. (B) The cumulative intensity distribution along NE contours standardized by subtraction of the mean and division by the standard deviation. Note that all Nups except Mpl1, Mlp2 and Ndc1 display highly consistent distribution patterns at the NE. (C) Intensity auto-correlation functions along the contours of NEs. Note that all Nups except Mpl1 and Mpl2 display very similar patterns of spatial variation. (D and E) NE intensities quantified with NuRIM in Nup84-yEGFP cells imaged (D) at different culture ODs and at the standard plating density or (E) at the standard OD and various plating densities. Mean ± SD; n - number of image frames analyzed. (F) Representative images illustrating co-localization of Gle1, Mlp1 and Mlp2 with Nup84 in the doubly labeled strains. Note that unlike Gle1, both Mpl1 and Mlp2 do not completely co-localize well with Nup84.

**Fig. S2.**
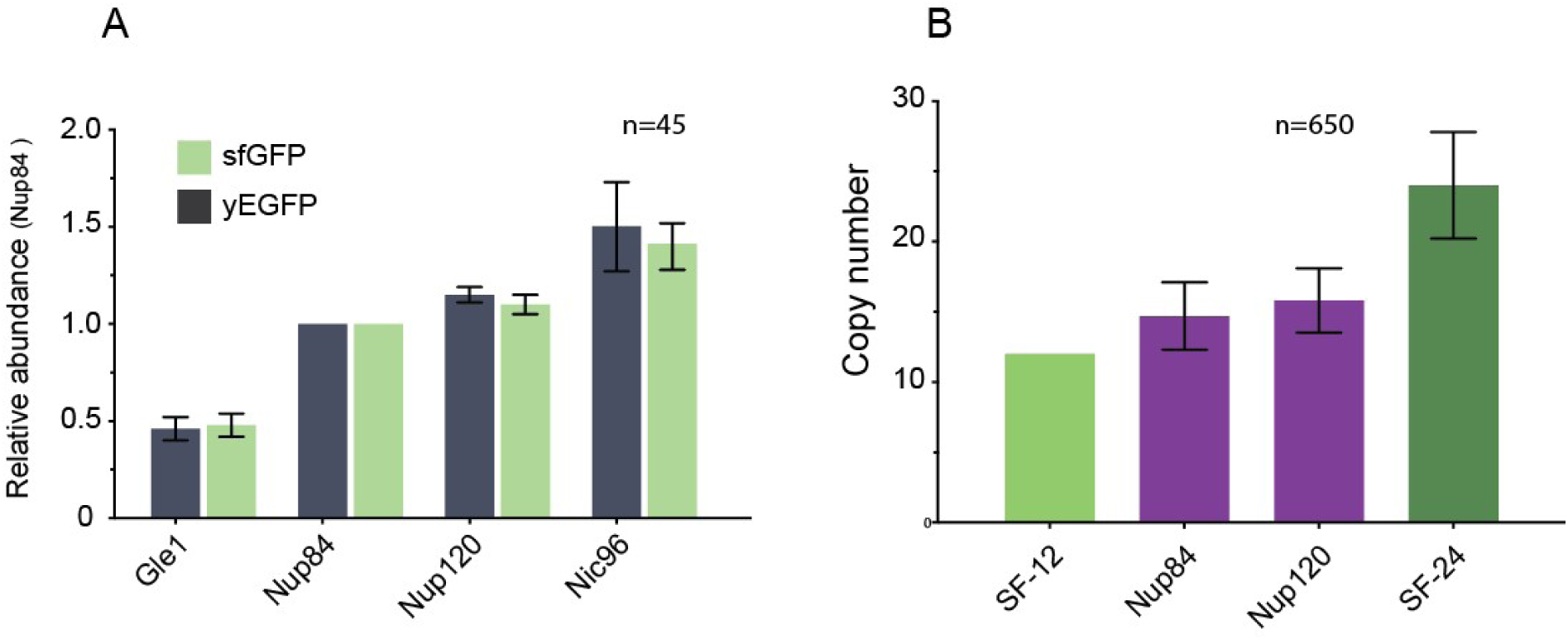
Analysis of ‘superfolder’ GFP tagged strains and absolute stoichiometry quantification for Nup84-sfGFP and Nup120-sfGFP. (A) The type of fluorescent tag does not affect relative stoichiometry values. NE intensities were quantified for the indicated Nups tagged with either yEGFP or sfGFP and normalized for the values obtained for Nup84. Mean ± SD; n - number of image frames analyzed. (B) The brightness of icosahedral particles carrying 12 and 24 sfGFP copies (SF12 and SF24) were compared to single NPC intensities in Nup84-sfGFP and Nup120-sfGFP expressing cells as outlined in Fig. 2 A and B. Mean ± SD; n - number of particles analyzed.

## 3) Supplementary tables

**Table S1.**
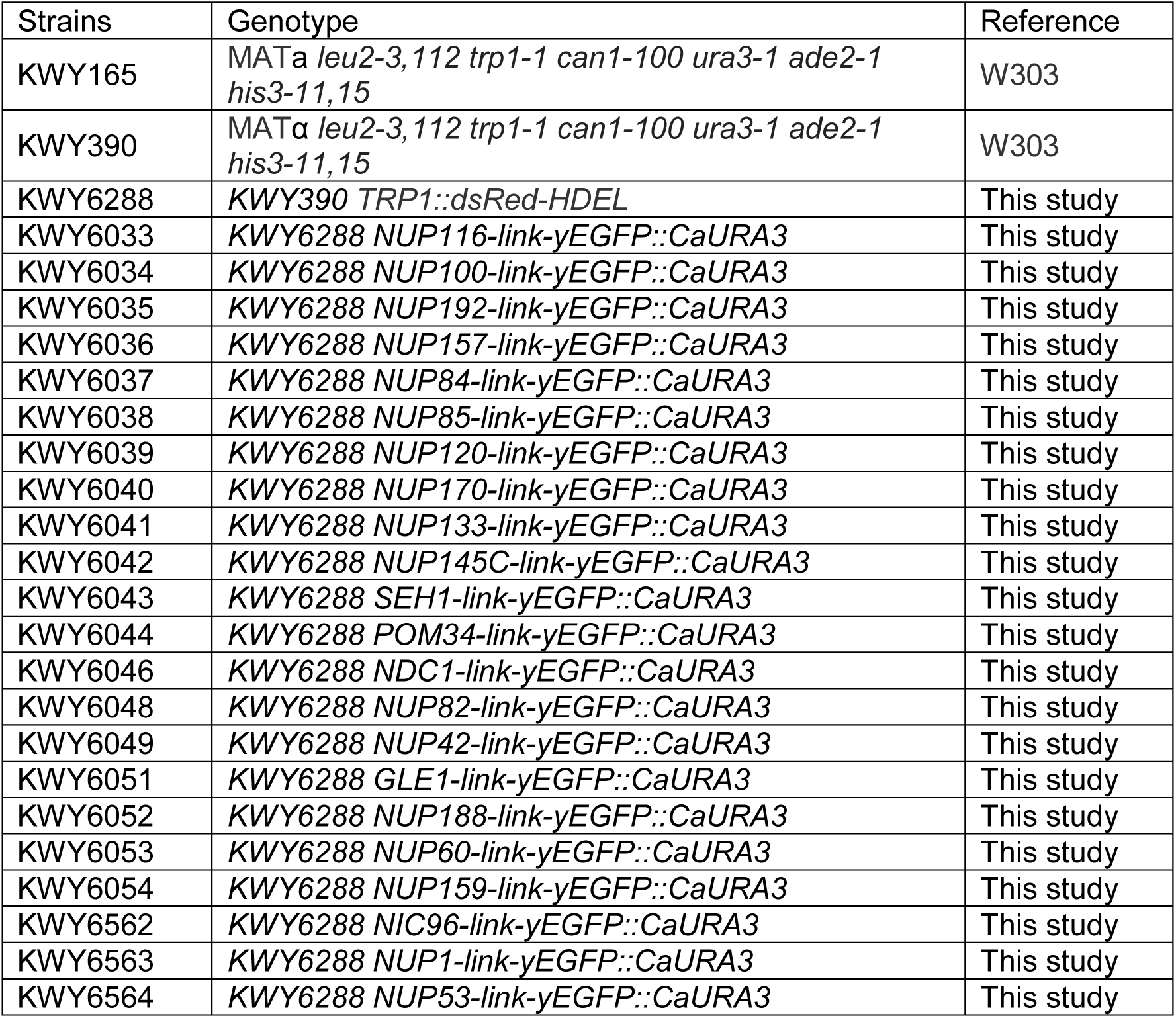
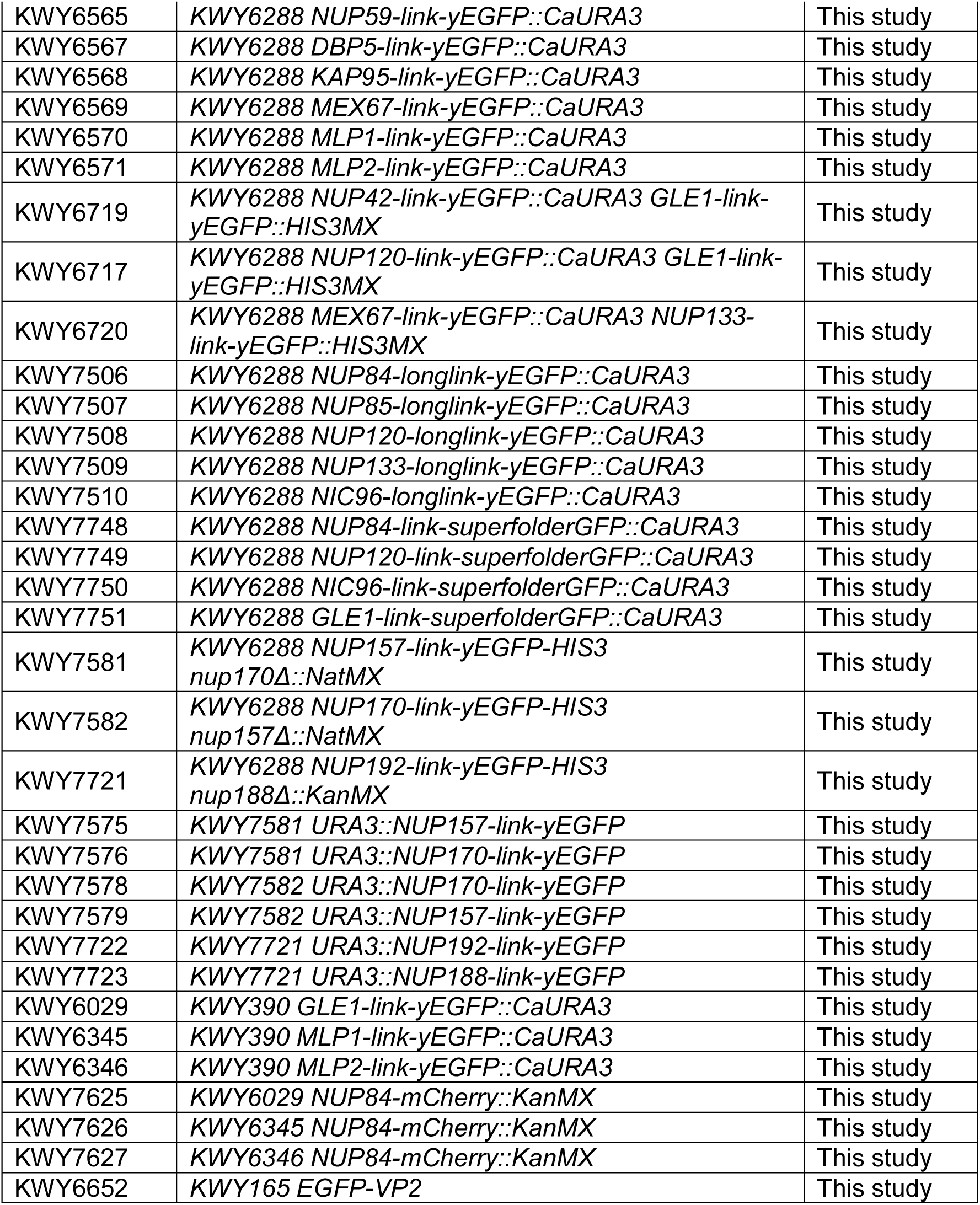
Strains used in this study

**Table S2.**
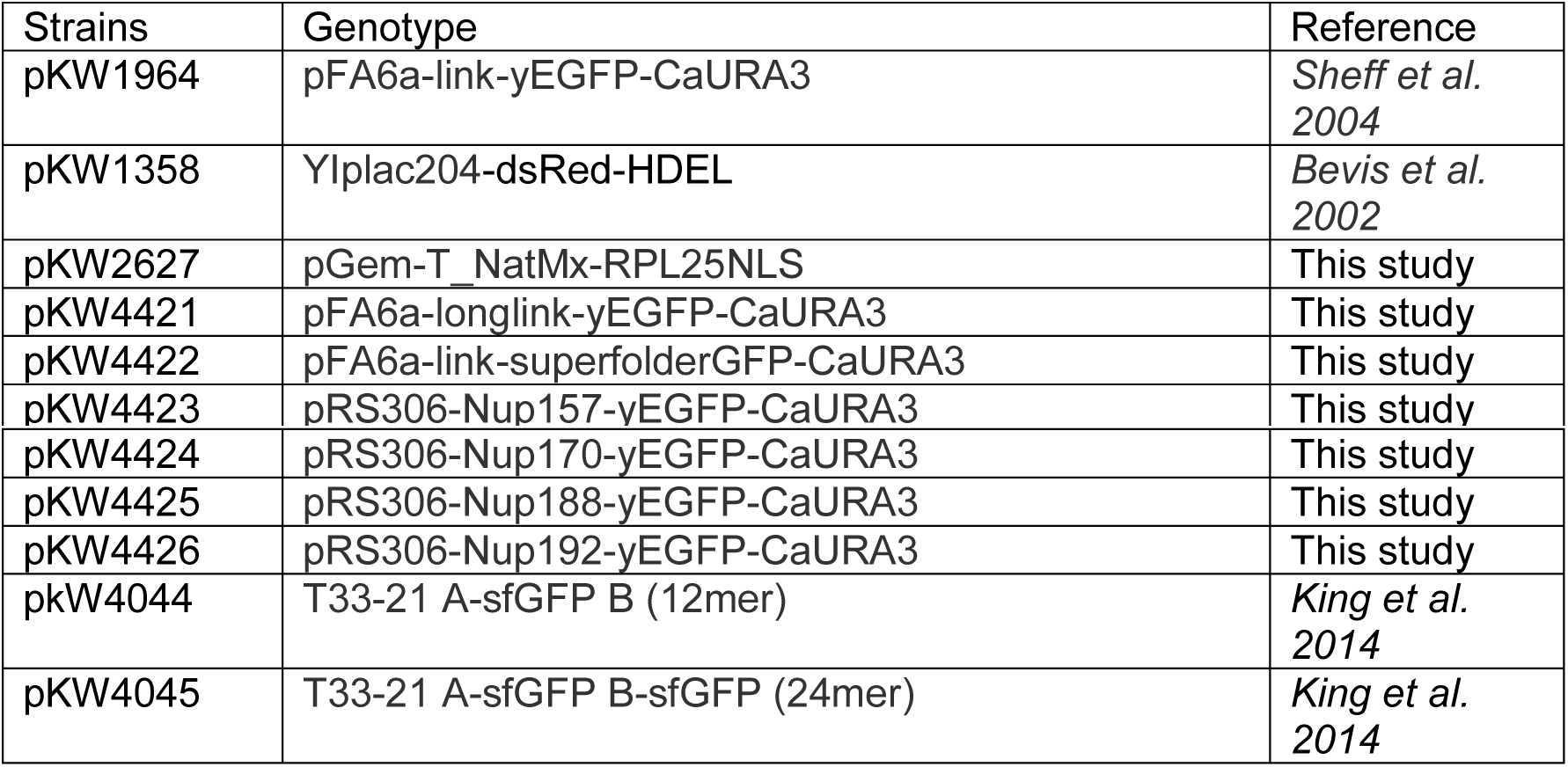
Plasmids and constructs used in this study.

## 4) Supplementary Movies

Movie S1: A Z-stack fluorescence image projection of dividing Nup84-yEGFP cells illustrating separate NPCs at the NE region that bridge the mother and daughter nuclei.

Movie S2: Single molecules of yEGFP were physisorbed on a glass coverslip. Comparing their fluorescence intensities with that of VP2 particles was compatible with 120 yEGFP molecules per VP2 particle.

